# NtrC expression at lag-phase improves competitive fitness under low and fluctuating nutrient condition

**DOI:** 10.1101/2022.03.29.486326

**Authors:** L K Mishra, R Shashidhar

## Abstract

The enteric pathogens cycle between nutrient rich host and nutrient poor external environment. These pathogens compete for nutrients in the host as well as in external environment and also often experience starvation. In this context, we have studied the role of a global nitrogen regulator (NtrC) of *Salmonella* Typhimurium. The *ntr*C mutation caused extended lag phase and slow growth in minimal media. In lag phase the wild-type cells showed ∼60 fold more expression of *ntr*C as compared to log phase cells. The role of *ntr*C gene is often studied with respect to nitrogen scavenging in a low nitrogen growing condition. However, our observation indicates that, even in the adequate supply of nitrogen, the *ntr*C null mutants failed to adapt to new nutrient conditions and were slow to exit from the lag phase. Gene expression studies at lag phase showed down-regulation of the genes involved in carbon/nitrogen transportation and metabolism in Δ*ntr*C mutant. In the co-culture competition studies, we observed that *ntr*C knockout was unable to survive with the wild-type *Salmonella* and *E*. coli. We also observed that the Δ*ntr*C mutant did not survive long term nitrogen starvation (150 days). Critical analysis of starvation survival revealed that, *ntr*C gene is essential for recycling of nutrients from the dead cells. The nutrient recycling efficiency of Δ*ntr*C mutant was ∼ 12 times less efficient than wild-type. Hence, the current work establishes that *ntr*C expression at the lag phase is essential for competitive fitness of *Salmonella* to survive in an environment having low and fluctuating nutrient conditions.

**Importance:** *Salmonella* Typhimurium, both inside or outside of the host organism face enormous competition from other microorganisms. The competition may take place either in steady or in fluctuating climatic conditions. Thus, how *Salmonella* Typhimurium survive under such overlapping stress conditions, remained unclear. Therefore, here we report that, a global regulator NtrC, which is also a part of two-component system, activates the set of genes and operons involved in rapid adaptation and efficient nutrient recycling/scavenging. These properties, make cell able to compete with other microbes, under characteristic feast-or-famine life style of *Salmonella* Typhimurium. Therefore, this piece of work helps us to understand the starvation physiology of enteric bacterial pathogen *Salmonella* Typhimurium.

## Introduction

The enteric bacterial pathogens *Salmonella* and *E*. coli often experience fluctuating nutrient conditions, as they cycle between host and external environment. In external environment, these enteric bacterial pathogens exposed to extreme conditions, including extensive competition among microbes for limited nutrients and ultimately nutrient starvation (1-4). To cope with nutrient starvation and competition, enteric bacteria have evolved comprehensive mechanisms to scavenge nutrient from environment (5-7). Among all the nutrients, after carbon, nitrogen (N) is the major macromolecule nutrient for the cell as it is the part of nucleic acid and protein. While, ammonia is most favoured nitrogen source(8). In nitrogen limited condition cell activates nitrogen scavenging system to assimilate N from alternative sources (9, 10). In *Salmonella* and *E*. coli the two component system, NtrBC, is activated during N limited conditions (11, 12). The NtrB (gene *gln*L) is the sensor with kinase activity and NtrC (gene *gln*G) is the DNA binding response regulator(12). During N starvation, NtrC is phosphorylated and activate multiple genes across the genome (13). There are many reports on the molecular mechanism of NtrBC and the genes regulated by this two component system(8). However, NtrC is not an essential protein and null mutants behave like wild-type except in nitrogen limited condition(14). Hence, most of the research has been carried out in this direction. However, there is only one report on the role of NtrC in survival of *Salmonella* in long term starvation (more than 30 days)(15). Also, there are no reports on the role of NtrC in competitive fitness. Therefore, taking clue from the earlier work on NtrC, we explored the role of NtrC in survival of *Salmonella* Typhimurium during long term starvation and competitive fitness in low and fluctuating nutrient condition in environment, by generating isogenic knockout mutant of *ntr*C (*gln*G) gene. Our results suggest that, NtrC not only facilitates rapid adaptation to the changing environments but also improve nutrient recycling efficiency of the cell. These two features together enhance the overall competitive fitness of *Salmonella* Typhimurium.

## Materials and methods

### Bacterial strains and growth conditions

The culture of *Salmonella enterica subsp. enterica* serovar Typhimurium strain LT2 (MTCC 98) was obtained from Microbial Type Culture Collection and Gene Bank (MTCC) Chandigarh, India. Luria broth, (LB) was used for the overnight growth of strains in all the experiments. M9 was used as a base for minimal medium (6g/L Na_2_PO_4_, 3g/L KH_2_PO_4_, 1g/L NH_4_Cl, 0.5g/L NaCl, 2mM MgSO_4_, 0.1mM CaCl_2_) and was supplemented with 0.4% glucose whenever needed. The cultures were incubated at 37°C with shaking at 120 rev/min. LB agar plates and those supplemented with kanamycin (50µg/ml) and carbenicillin (50µg/ml) were used wherever required.

### Gene knock-out mutant preparation

The *ntrC (glnG)* gene knock-out mutant (Δ*ntrC* mutant) was generated by using Quick and Easy *E. coli* gene deletion kit by Red^®^/ET^®^ Recombination, Gene Bridges, (Heidelberg, Germany). The sequence for *ntrC* gene of *S*. Typhimurium was obtained from NCBI (NC_003197.2). The mutant was confirmed phenotypically, through newly acquired antibiotic (kanamycin) resistance. Genotypically, by diagnostic PCR. The mutant characteristic phenotypes, was matched with the phenotype given in literatures. The pJET1.2/blunt cloning system (ThermoFisher Scientific, Massachusetts, USA) was used for complementation of *ntrC* gene with its native promoter (pntrBC *glnLG*). The primers and their sequences used in the study are given in supplementary S1.

### Growth studies in different nutrient medium

To study the growth curve of wild-type and Δ*ntr*C mutant,198µl of fresh LB or M9 (supplemented with 0.4% glucose) was taken in 96 well microtiter plates along with 2µl of overnight grown culture. Kanamycin was added in mutant culture (50µg/ml) and the plate was placed in BioTek Synergy H1 plate reader (Winooski, Germany). The reading was taken for 24 hours in form of O.D at 600nm, with continuous shaking condition at 37°C.

### Survival studies in different nutrition deficient conditions

The wild-type and the Δ*ntr*C mutant (100µl) was inoculated in 10ml M9 (1x) minimal media with complete and nutrient limited conditions (without carbon, nitrogen and phosphorus). At regular intervals cell survival was evaluated using the standard plate count method.

### Competition studies in batch culture

For competition assay, 1ml of both wild-type and **Δ***ntrC* mutant was co-inoculated in 50ml fresh M9 media supplemented with 0.4% glucose (∼10^8^ cells/ml). The competition assay was conducted in 250ml conical flask under sterile condition. The competition was continued till four months without any further addition of nutrient (Fig.3a). The cell number was calculated after every month by dilution and plating method.

### Competition studies under feast-or-famine cycle

The culture containing both wild-type and Δ*ntr*C mutant (50µl) of one-month competing cells from above running experiments (competition in batch culture) was taken in 50ml fresh M9 media. Competition was conducted in 250ml conical flask under sterile condition. The competition was conducted for one month. After one month of competition 100µl of culture were transferred to 50ml fresh M9 medium. The cycle of competition for one month and then replenishment with fresh media was repeated four time(Fig.3c). The cell count was taken after 24 hours of incubation in fresh medium.

### Nutrient recycling efficiency of starved cells

To study the nutrient recycling efficiency of the starved cells, the protocol was followed as described by Schink *et al*. (2019) with some modifications in it(16). Some fraction (15ml) of starved cells was taken, centrifuged at 5000 rpm for 10 minute. Re-suspended in equal volume of sterile distilled water and transferred into sterile glass petri plate for exposure to UV light for 30 minute from the distance of 5cm (230 V, 50/60 Hz) VILBOUR LOURMAT (Collegien, France). The UV killed cells used as growth medium for starved cell (Fig.4b). The viable starved cell (10µl) was inoculated in 990µl dead cell solution in 1:100 ratios for two days. The growth/survival of starved cell was analysed after 2 days, by dilution and plating method (CFU/ml). The efficiency of nutrient recycling was determined as described in Schink et al (2019). The equation used to quantify the nutrient recycling efficiency is given in Fig.4c.

### Gene expression studies

For gene expression studies, the total RNA extraction was carried out by the TRIzol method using TRIzol reagent (Invitrogen), and cDNA synthesis were carried out using TransGen (Beijing, China) reagent and protocol. The primers were designed using ‘Integrated DNA Technologies Primer Quest software’ (www.idtdna.com/site). The 16s rRNA was used as the housekeeping gene. The cDNA products were subjected to SYBR green RT-qPCR assay using primers and DyNAmo Flash SYBR Green qPCR Kit Finnzymes (Espoo, Finland) in amplification master cycler, Roche (Basel, Switzerland) at 94°C for 10min, followed by 40 cycles consisting of denaturation at 95°C for 10 s, annealing at 55°C for 10 s and extension at 72°C for 20 s. Following amplification, determination of threshold cycle (CT) values and melting curve analysis was carried out. The analysis was carried out following the MIQE guidelines for real-time PCR experiments(17).

### Study of competition assay using reporter gene

The wild-type and Δ*ntr*C mutant was transform with plasmid, pDsRED expression vector system, containing Red fluorescent protein (RFP) and under the lac promoter. Similarly, another pDs vector system was used in which instead of RFP, green fluorescent protein (GFP) was used as a reporter gene. The cells transformed with RFP or GFP vector (pDs vector containing either GFP or RFP) were selected on Ampicillin (100-µg/ml). The transformation was confirmed by diagnostic PCR and observation of transformed cells under the fluorescent microscope. The competition assay was employed in 96 well microtiter black colour plates with transparent cover at the top in BioTek Synergy H1 plate reader (Waldbronn, Germany). The GFP excitation/emission was monitored at 395nm/509nm while RFP fluorescence was monitored at excitation/emission was monitored at 558nm/583nm

### Glucose estimation assay

For glucose estimation, 10^7^ cells of washed wild-type and Δ*ntr*C mutant were incubated in 1ml M9 minimal media, supplemented with 1mg/ml of glucose. At different time points the unused glucose in the minimal media was estimated using Sigma Glucose (GAGO20) assay kit (New jersey, USA). To measure glucose concentration in solution, we followed the instruction given in the kit.

### Statistical analysis

All the experiments were carried out in duplicates and were repeated three times. In all cases, population was tested as a random factor and environmental regime was tested as a fixed factor. Student’s t-tests were performed in Microsoft Excel. Significance was first assessed using α = 0.05 as the cut-off (p <0.05).

## Results

### *ntr*C mutation caused extended lag phase of *S*. Typhimurium in minimal medium

The NtrC is an important nitrogen metabolism regulator. Therefore, the effect of *ntr*C null mutation on the growth of *S*. Typhimurium in various media was evaluated. In the protein rich medium (Luria broth), wild-type and Δ*ntr*C mutant showed no significant difference in the growth (Fig.1a). However, in M9, Δ*ntr*C mutant showed extended lag phase and slow growth rate, as compared to wild-type. The Δ*ntr*C mutant has lag phase of 8-hr and growth rate of 48-minutes. While wild-type has lag phase of 3-hr and growth rate of 44-minutes (Fig.1b). The Δ*ntr*C complement strain rescued the extended lag-phase phenotype (supplementary S2).

**Fig.1:**
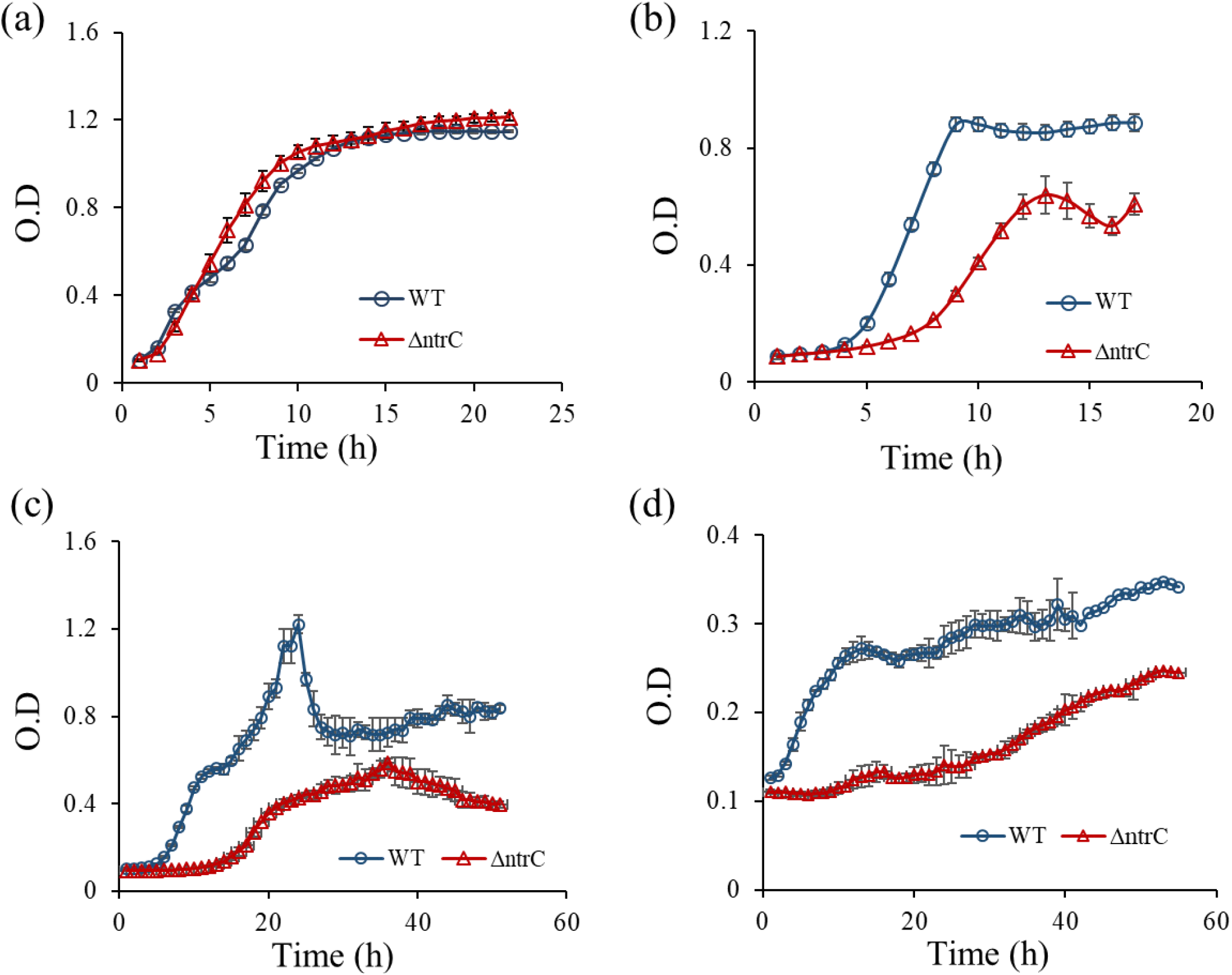
Growth characteristics of wild-type and Δ*ntr*C mutant. (a) In nutrient rich Luria broth medium. (b) In M9 minimal media supplemented with 0.4% (w/v) glucose. (c) The growth response of one month starved, wild-type and Δ*ntr*C mutant, in fresh M9 medium. (d). The growth response of one month starved, wild-type and Δ*ntr*C mutant, in dead cell extract.

### The *ntr*C is needed for faster recovery from long term starvation

The lag phase has been associated with the adaptation to the external environmental conditions(18). Further, it is often found that the lag phase of the stressed cell is longer than the fresh cells(19, 20). Therefore, we studied the recovery of wild-type and Δ*ntr*C mutant from starvation. In starvation condition, often cells recycle the nutrients released from the dead cells. Hence, recovery of starved cells was studied in both fresh M9 medium and dead cell extract as the nutrient medium. The wild-type cells showed rapid adaptation to newly provided condition, while Δ*ntr*C showed delayed response to this change, consequently, in M9 medium the lag phase of Δ*ntr*C mutant was 14-hrs long. The wild-type cells attained maximum growth (O.D ∼1.2) within 24-hrs of exposure to fresh media, whereas Δ*ntr*C mutant took 36-hrs to attained its maximum growth (O.D ∼0.6) (Fig.1c). Wherein dead cell extract was used as growth medium, the wild-type was able to grow faster than Δ*ntr*C mutant (Fig.1b). These results indicate that *ntr*C is required for rapid adaptation to environmental changes.

### *ntr*C knockout of *S*. Typhimurium unable to survive under long term nitrogen starvation

The extended lag phase and slow recovery from starvation prompted us to study the effect of *ntr*C mutation on long term nutrient starvation tolerance. Often microorganisms with slow growth or extended lag phase exhibit better survival under starvation (21, 22). Therefore, Δ*ntr*C mutant was examined for its survival under nutrient limitation. The wild-type and Δ*ntr*C mutant was grown in M9 media, which is devoid of any one of these macro nutrient (carbon, nitrogen or phosphorous). The survival of both strains in nutrient limiting medium was compared with complete M9 media. The wild-type and Δ*ntr*C mutant maintained their cell number, ∼ 10^7^ cells/ml, in complete M9, in carbon and phosphorous limited M9 medium (Fig.2(a-c)) respectively. However, in nitrogen limited M9 medium, the Δ*ntr*C mutant showed complete death within 150-days, while wild-type was able to maintain its survival and attained stable cell number,∼ 10^4^ cells/ml (Fig.2d).

**Fig.2:**
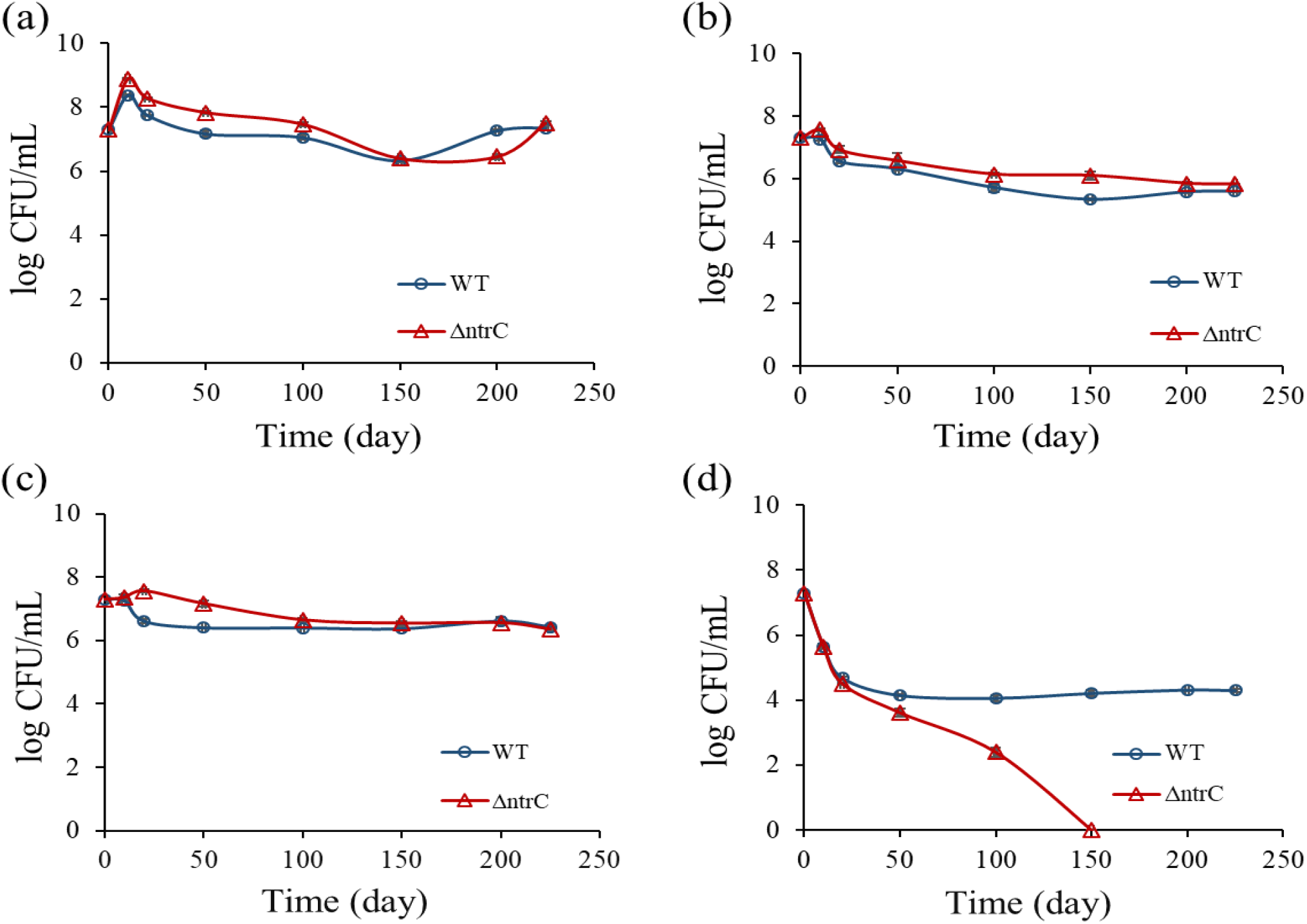
Survival studies of wild-type and Δ*ntr*C mutant in different nutrient limitations. (a) In M9 complete medium (b) In M9-C (carbon limited) medium (c) In M9-P (phosphorous limited) medium (d) In M9-N (nitrogen limited) medium.

### *ntr*C null mutant of *S*. Typhimurium is a poor competitor in constant and fluctuating nutrient condition

Next, the Δ*ntr*C mutant fitness during competition for nutrient utilization was evaluated. The wild-type and Δ*ntr*C mutant were incubated together (co-culture) in M9 medium for up to 4 months(Fig.3a). At the end of 4-months of incubation in M9 medium, wild-type cell number reduced by 99% (2-log reduction) and Δ*ntr*C numbers reduced by 99.9 % (3-log reduction) as compared to zero-day cell numbers (Fig.3b). The results show that Δ*ntr*C mutant population is gradually reducing, in competition with wild-type.

**Fig.3:**
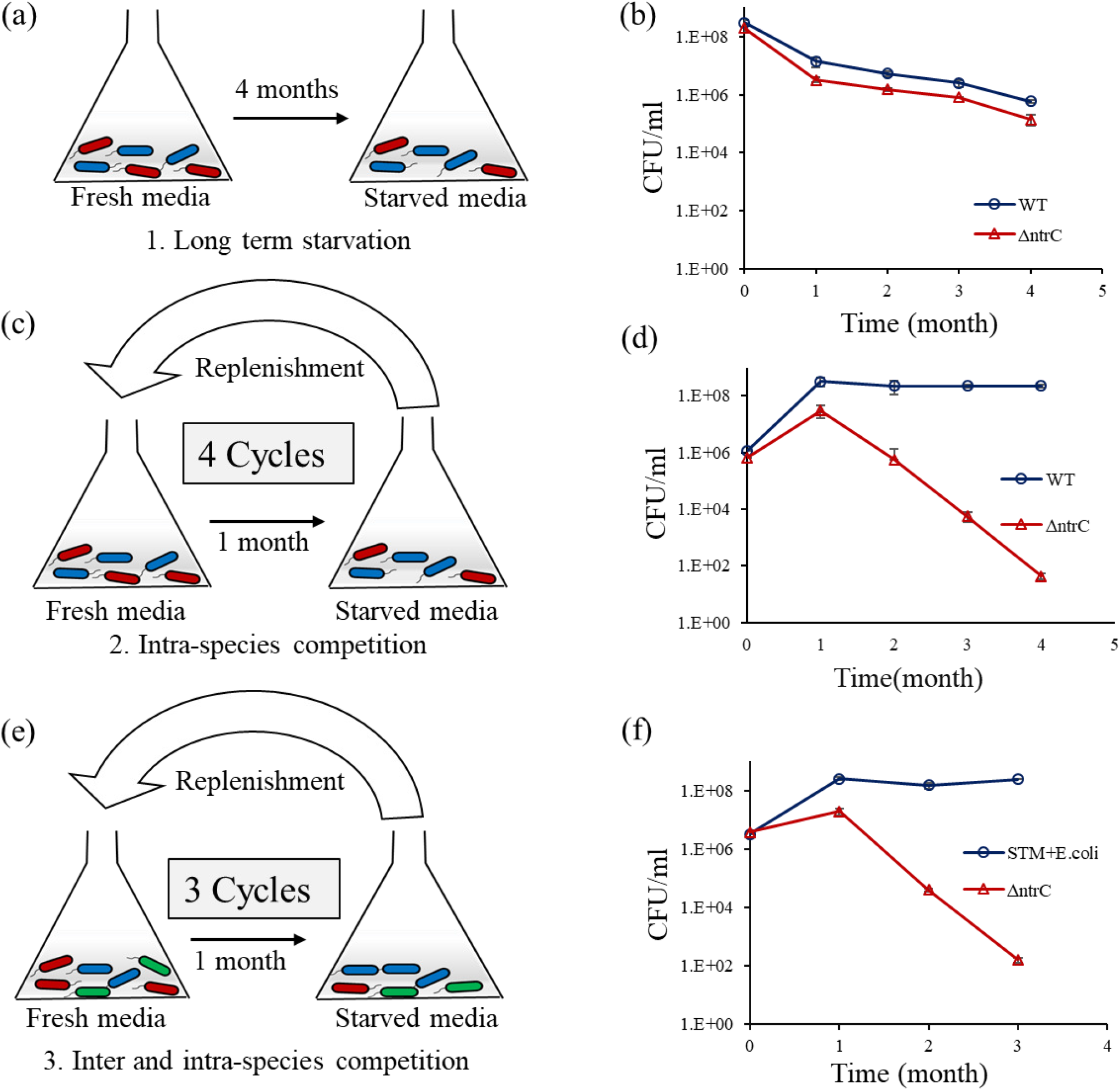
Study of the competitive fitness of Δ*ntr*C mutant in different conditions. (a) Schematic diagram showing competition under long term nutrient starvation (in batch culture). (b) Competition in batch culture, Δ*ntr*C mutant population is declining at relatively faster rate than wild-type strain. (c) Schematic diagram of competition under serial transfer to fresh medium (intra-species competition with wild-type cells). (d) Competition under serial transfer to fresh medium, shows that Δ*ntr*C mutant population is declining rapidly. (e) Schematic representation of inter and intra-species competition under serial transfer to fresh medium (f). The Δ*ntr*C mutant competition with wild-type *S*. Typhimurium and *E*. coli, showed even more rapid reduction after every transfer to fresh medium.

In another set of experiment, the competing cells (wild-type and Δ*ntr*C mutant) were transferred to fresh M9 medium every month(Fig.3c). The Δ*ntr*C mutant showed significant reduction in cell number after each cycle. At the end of 4 cycles of replenishment with fresh medium, the Δ*ntr*C mutant cell number reduced by 6-log, whereas, the wild-type cell number remained the same (Fig.3d). Further, similar nutrient replenishment experiment was carried out by adding one more competing species i.e. *E coli* (Fig.3e). After addition of *E. coli*, to the co-culture, the death of Δ*ntr*C mutant was accelerated, the Δ*ntr*C mutant population showed 6-log reduction within 3 cycles, instead of 4 cycles of nutrient replenishment. Therefore, increase in number of competing species accelerates the rate of reduction of Δ*ntr*C mutant population (Fig.3f).

### *ntr*C null mutant showed poor nutrient recycling efficiency

Starving microbial population feed on the nutrient released from the dying cells (Fig.4a). Hence, role of *ntrC* on nutrient recycling ability of *S*. Typhimurium was evaluated. The UV killed cell extract was used as growth medium for the viable cells (Fig.4b). The calculation used to measure the nutrient recycling efficiency (growth yield) is given in the Fig.4c. After 24-hrs of incubation in dead cell extract, the wild-type showed efficient nutrient recycling (absolute growth yield of viable cell on dead cell), α= 0.185, in unit of “new cell” per “killed cell”. Hence, nutrients released from ∼5.4 cells are required to regenerate one new cell. Whereas, the Δ*ntr*C mutant showed very poor efficiency of nutrient recycling, α= 0.015 which means Δ*ntr*C mutant need ∼ 66 cell to make one new cell (16). The result shows that Δ*ntr*C mutant has almost 12 times less nutrient recycling efficiency than the wild type *S*. Typhimurium. Since, NtrC is an activator of nitrogen transporter gene. So, nutrient recycling efficiency in Δ*ntr*C mutant may be limited by nitrogen. Therefore, we supplemented dead cell extract with 10mM ammonium chloride and analysed nutrient recycling efficiency. There was no improvement in the growth of Δ*ntr*C mutant (supplementary S2). Hence, we conclude that the poor nutrient recycling efficiency of Δ*ntr*C mutant, is not nitrogen limiting.

**Fig.4.**
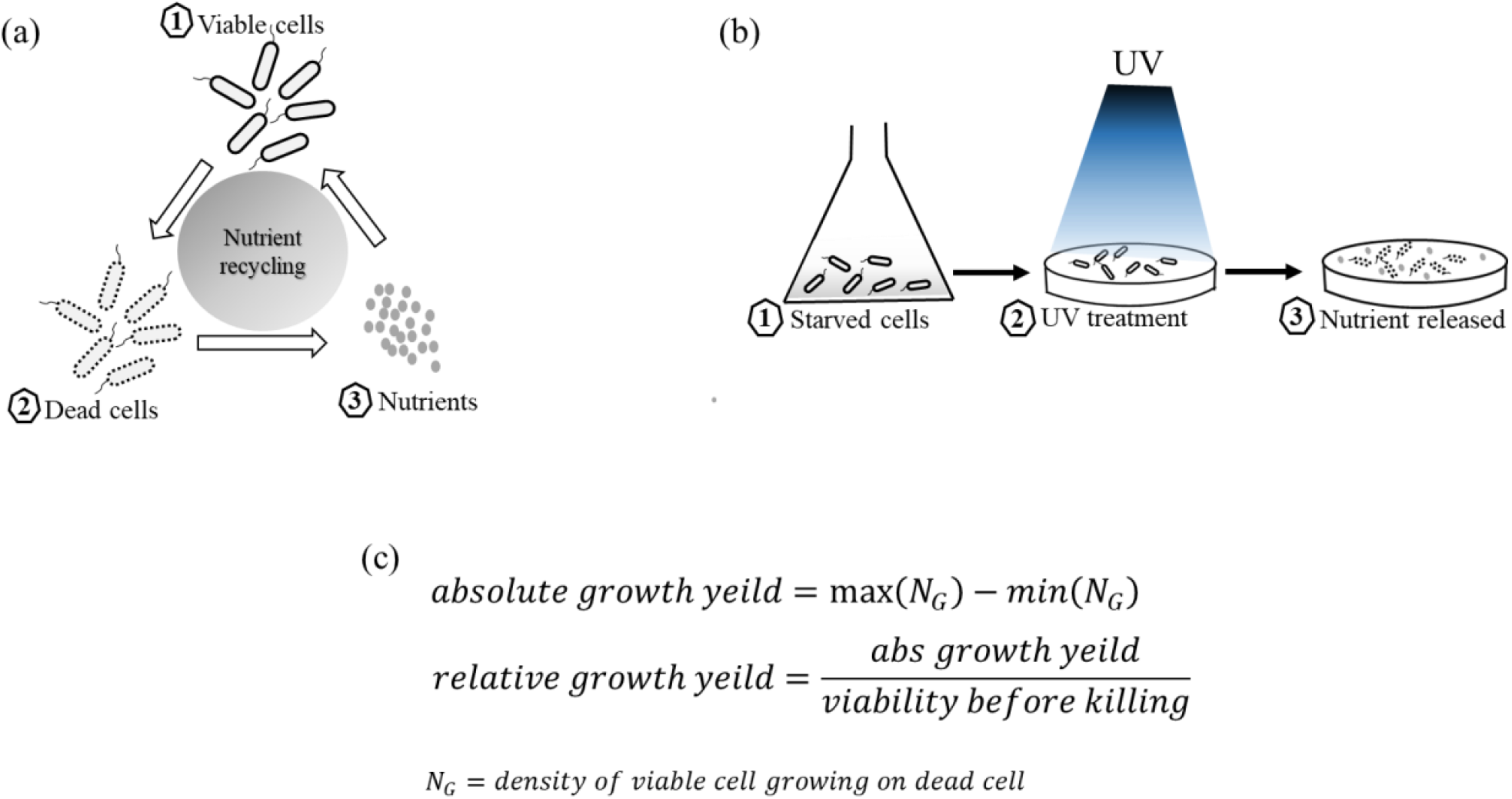
Nutrient recycling efficiency: (a) Schematic representation of the process of nutrient recycling by the bacterial cells during nutrient starvation. (b) Procedure used to kill the starved cells, for more detail see the “material and methods”. (c) Formula used to calculate the nutrient recycling efficiency.

### Slow growth and poor nutrient recycling efficiency eliminates Δ*ntr*C mutant from competition

In order to validate the results obtained from the last two experiments, slow growth response and poor nutrient recycling efficiency of Δ*ntr*C mutant. The, 10^8^ cells/ml, of wild-type and Δ*ntr*C mutant was employed for competition in distilled water (23-25). Here, the nutrient released from the dying cells, are the only source of nutrient for the competing cells. Therefore, the cell equipped with rapid and highly efficient nutrient scavenging system will survive and grow. When wild-type and Δ*ntr*C mutant were incubated separately, without competition (Fig.5a), wild-type population stabilised at ∼10^4^ cells/ml and Δ*ntr*C mutant population was completely eliminated after two-month of starvation in distilled water (Fig.5b). However, when wild-type and Δ*ntr*C mutant were kept together (co-inoculated) for competition in distilled water (Fig.5c), the Δ*ntr*C mutant survival remained unaffected consequently died after two months, but wild-type cell number stabilized at, ∼10^6^ cells/ml (Fig.5d). The 2-log (99%) increases in wild-type cell number, suggest that wild-type is quickly utilising the entire nutrient released from the death of both type of cells. For further confirmation, the competitive fitness of starved cells of both strains were observed in real time, using reporter genes. Competition in M9 media, showed that the wild-type was growing normally while Δ*ntr*C mutant grow slowly (Fig.6a). Similarly, during competition in dead cell extract, the Δ*ntr*C mutant again showed less growth than wild-type (Fig.6b).

**Fig.5.**
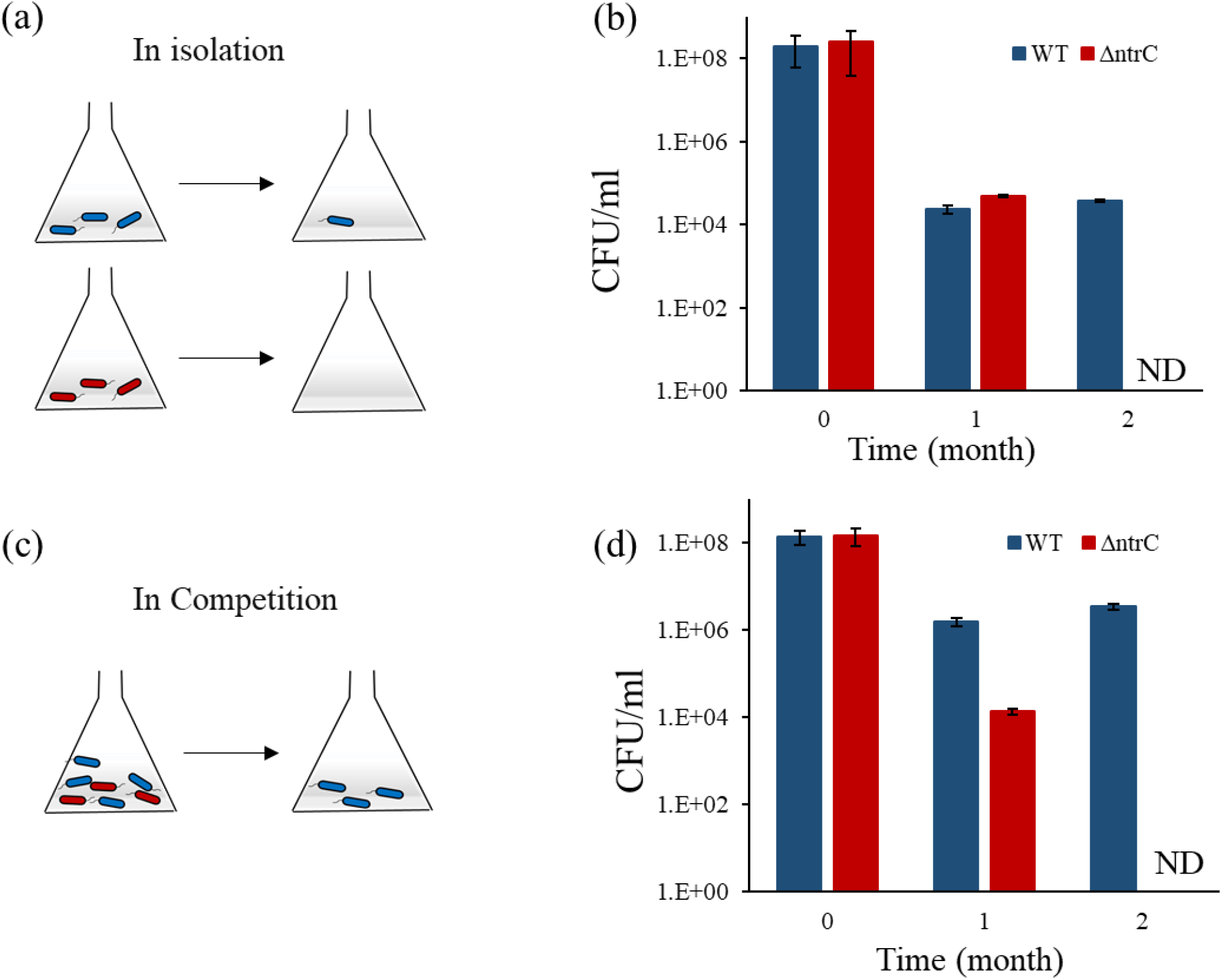
Survival of wild-type and Δ*ntr*C mutant in nutrient free medium:(a) Survival of wild-type and Δ*ntr*C mutant in distilled water, without competition. (b) Survival of wild-type and Δ*ntr*C mutant in distilled water, with competition. ND indicates that no viable cells of Δ*ntr*C mutant was found (not detected).

**Fig.6:**
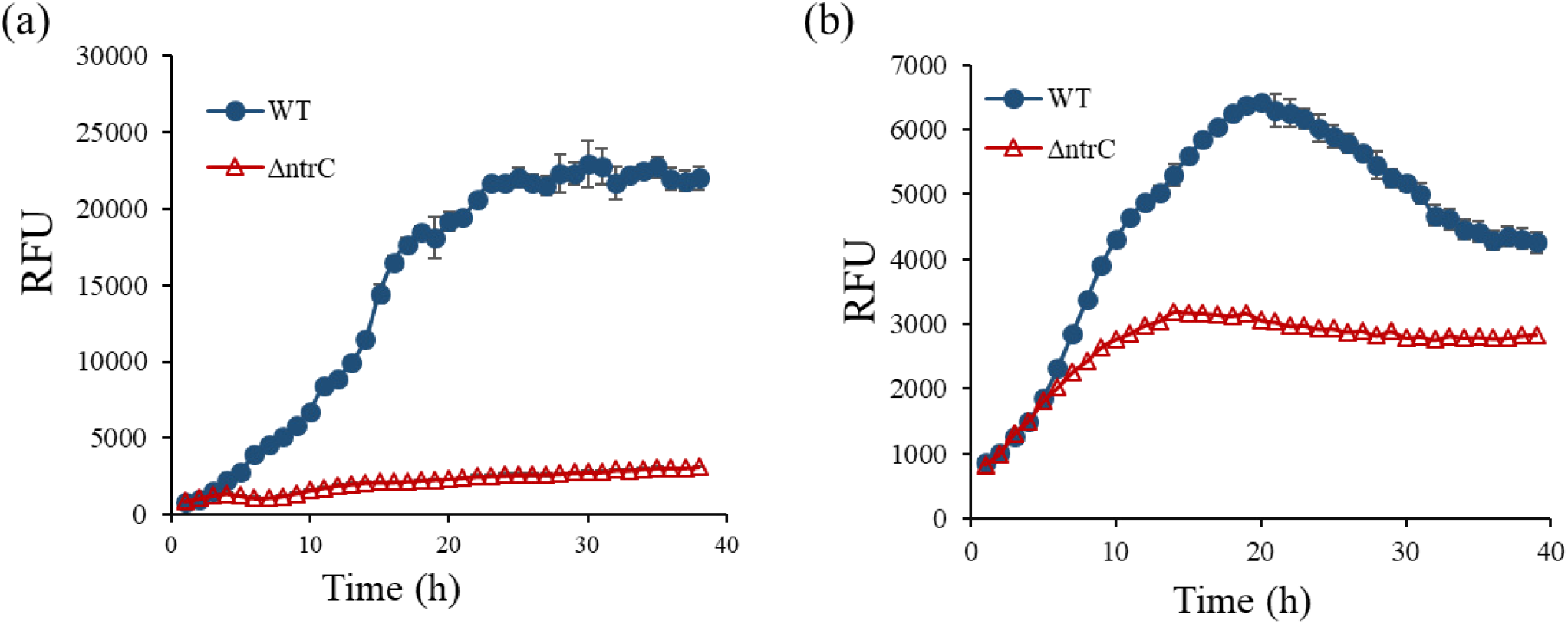
Real-time analysis of competitive fitness of wild-type and Δ*ntr*C mutant: the competition was monitored by using reporter gene GFP (wild-type) and RFP (Δ*ntr*C mutant). (a) Competition in fresh M9 medium. (b) Competition in dead cell extract. The wild-type strain showed more efficient growth than Δ*ntr*C mutant.

### *ntr*C is required in early phase of growth of *S*. Typhimurium in minimal medium

The growth, fitness and nutrient recycling experiments prompted us to study the role of *ntr*C gene in growth at early phase (lag phase) in minimal media. Thus, the important genes involved in the nitrogen and carbon metabolism were evaluated in Δ*ntr*C mutant.

In first set of gene expression studies, the wild-type cells were analysed for expression of *ntr*C gene at different stages of the growth. Interestingly, in lag phase we observed 60-fold expression with respect to expression at 6-hr (Fig.7a). In second set of experiment, we analysed the expression of 22 genes involved in both carbon and nitrogen transport and metabolism in lag phase of *ntr*C knockout strains with respect to wild-type. The genes involved in glucose transport (*pts*G, *Pts*H, *crr*,) were 4 to 6 fold down-regulated. The *amt*B, encoding ammonium transporter, showed ∼32 fold down-regulation. The gene involved in nitrogen metabolism *gln*A, *gdh*A, *arc*B, *fix*A, *ast*C, *ast*A showed 7-10 fold down-regulation, while *arc*A, *lys*A, *gla*H showed 5-7 fold down-regulation. The glucose metabolism gene *sdh*A, *agp, glt*A, *STM0761* showed 3 to 4 -fold down-regulation. While, gene *ace*B and *STM2757* showed 5 to 7 fold down-regulation (Fig.7b). The result indicates that Δ*ntr*C has overall slow metabolism than the wild-type *S*. Typhimurium. To evaluate the result obtained from gene expression studies, we compared the glucose uptake by wild-type and Δ*ntr*C mutant. The wild-type cells utilized glucose at much faster rate than Δ*ntr*C mutant. The wild-type cells consumed glucose within 5-hrs, while Δ*ntr*C mutant took 8-hrs to consume the glucose present in the growth medium (Fig.7c).

**Fig.7:**
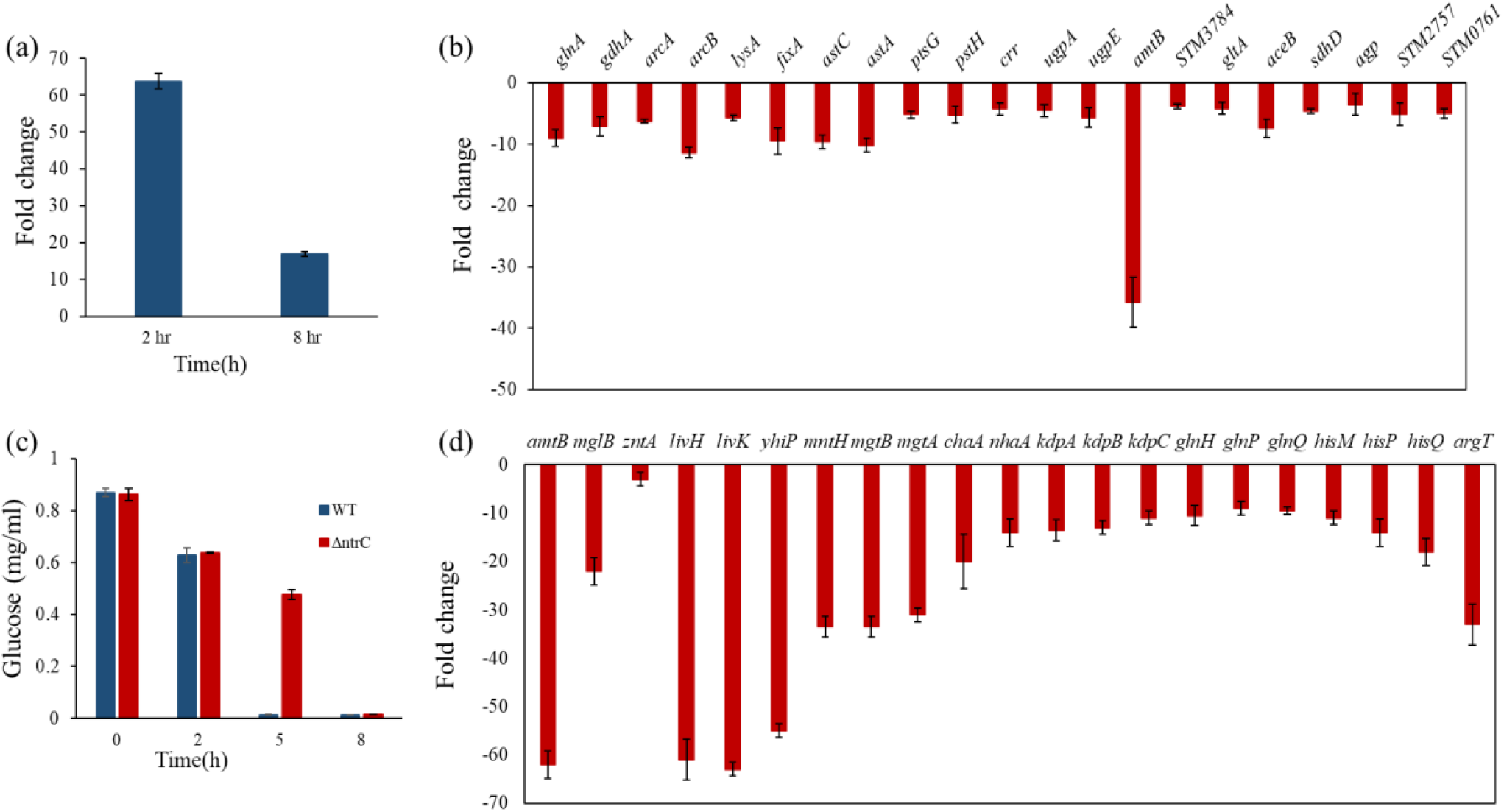
Molecular basis of slow adaptation and growth:(a) The *ntr*C gene expression in wild-type strain.(b) Nitrogen/carbon transport and metabolism genes expression analysis in Δ*ntr*C mutant. (c) Glucose estimation in wild-type and Δ*ntr*C mutant. (d) Gene expression study for nutirent recycling efficiency.

In the third set of gene expression studies, our objective was to understand the reason for poor nutrient recycling efficiency of Δ*ntr*C mutant. Hence, we studied the genes which are involved in the scavenging of nutrients from the environment. After two hour of incubation in dead cell extract, the genes of Δ*ntr*C mutant was evaluated with respect to wild-type. The genes involved in the utilization of nitrogen containing compounds like ammonia, amino acids, peptides, in Δ*ntr*C mutant were down-regulated. Genes involved in ammonia transport *amt*B, was ∼62 fold down-regulated while the gene coding for dipeptide and tripeptide permease, *yhi*P, was ∼55 fold down-regulated. Similarly, gene coding for high affinity branched amino-acid ABC transporter permease, *liv*K and *liv*H was ∼60 fold down-regulated, genes coding for transport and utilization of amino-acids like glutamine (*gln*H, *gln*P, *gln*Q) histidine (*his*M, *his*P, *his*Q) and arginine (*arg*T) was 10-20 fold down-regulated (Fig 7d). Moreover, we also observed the expression of the genes coding for transporter proteins of metal ions, like magnesium, sodium, potassium, zinc (*mgt*A, *mgt*B, *mnt*H, *nha*A, *kdp*A, *kdp*B, *kdp*C) all of these genes were down-regulated in Δ*ntr*C mutant, as compare to the wild-type *S*. Typhimurium (Fig.7d).

## Discussion

The “Two-component signal” transduction systems enable bacteria to adapt the fluctuating environment(26). NtrB/NtrC is a two component signal transduction system, known to modulate multiple operon during nitrogen limited conditions(27). However, role of NtrC other than nitrogen regulation is very limited. Therefore, we have studied the role of NtrC in bacterial growth, long term starvation tolerance, and during competitive fitness.

### The NtrC is needed for faster lag phase exit in minimal medium

We noticed that mutation of *ntr*C caused slow growth in M9-minimal medium. Also, the *ntr*C mutation impaired the lag phase of M9 grown *S*. Typhimurium (Fig.1b). Therefore, we investigated the role of NtrC in bacterial growth during lag phase. We started our study by complementing the Δ*ntr*C mutant. The Δ*ntr*C complemented strain successfully rescued the extended lag phase phenotype. Besides this, the wild-type gene expression analysis showed the high expression of *ntr*C gene at lag phase. Apart from us, Rolfe *et al*. also observed up-regulation of *ntr*C (*gln*G) gene in lag phase of LB grown *S*. Typhimurium(28). These results confirmed NtrC is required during the early phase of bacteria growth, in minimal medium. The impaired lag phase of Δ*ntr*C mutant prompted us to study molecular mechanism. Since, ammonia, a nitrogen source in M9, at higher concentration is not growth limiting. The passive transport of ammonia is sufficient to support optimal growth(29, 30).Therefore, the expression of nitrogen metabolism genes in Δ*ntr*C mutant were analysed at lag phase (2-hrs). All the tested genes were down-regulated. Further analysis revealed that the glycolysis and TCA cycle genes were also down-regulated. The RT-PCR results were further supported by our observation on slow glucose consumption by Δ*ntr*C mutant. Hence, *ntr*C mutation hampers the process of gene activation, which in turn slows down the overall metabolism of *S*. Typhimurium. Thus, the extended lag phase in Δ*ntr*C mutant, indicates the role of NtrC protein in lag phase metabolism. The earlier studies on *ntr*C, showed that NtrC is required to activate the expression of genes involved in nitrogen utilization under low nitrogen condition(5, 9, 15). However, the role of NtrC in a normal growth conditions remains unexplored. Therefore, we report that, besides low nitrogen conditions, NtrC is crucial for early phase (lag phase) growth in minimal medium. Hence, on arrival of fresh nutrients, NtrC at lag phase, enhance the expression of genes involved in nutrient transport and metabolism. Which in turn, uptake and metabolize nutrients at faster rate, that help cells to exit lag-phase in short duration.

### The role of NtrC in long term starvation

Lag phase of bacterial growth has been associated with adaptation(18). Moreover, previous studies on *ntr*C in *E*. coli, showed interplay between NtrC and adaptation to nitrogen starvation. These studies were conducted in short term nitrogen starvation, from 20 minutes to 34-days (15, 31, 32). But, there is no study reported on the survival of Δ*ntr*C knockout mutant in long term nutrient starvation. Therefore, we investigated the role of NtrC in tolerance and adaptation to long-term starvation of carbon, nitrogen and phosphorus nutrients. Among all the tested conditions, Δ*ntr*C mutant could not tolerate nitrogen limitation for a long duration (150 day). For survive under long term starvation, cell adapt few strategies, including up-regulation of transcriptional regulator and subsequently transporter proteins, recycling of intracellular proteins, and recycling of nutrient from the dead cells(15, 25, 33). Warsi *et al*. have shown that within 34 days of growth in nitrogen limiting condition, the wild-type *E*. coli cells acquire mutations at *gln*L and *gln*G gene. These mutations result in up-regulation of expression of *ntr*C (*gln*G) gene. The up-regulated *ntr*C further enhance the expression of genes encoding transporter proteins(15). However, in a *ntr*C knocked-out mutant, the essential process of gene activation to scavenge and utilize the nutrients present in low concentration is lacking. Consequently, Δ*ntr*C knockout mutant died after 150 days of nitrogen starvation. Even, Switzer *et al*. have shown that, inhibition of transcription at the onset nitrogen starvation has detrimental effect nitrogen starvation survival ability of *E*. coli(31). Therefore, NtrC which is a transcriptional activator of nitrogen transporter and metabolism genes, is essential for cell survival in long term nitrogen starvation.

### NtrC conferred competitive fitness to the *S*. Typhimurium in a stable and fluctuating nutrient conditions

Since, the length of lag phase experienced by the starved cells have been associated with the competitive fitness(20). Therefore, the role of *ntrC* in competitive fitness was evaluated. We observed that the Δ*ntr*C mutant failed to survive long term nutrient starvation and feast-or-famine condition, in a co-culture setup with wild-type cells. The Δ*ntr*C mutant gradually reduced in long term starvation, while in feast-or-famine condition showed rapid reduction in its population. Detailed analysis revealed that suppressed expression of nutrient scavenging and metabolism genes at lag phase, made Δ*ntr*C mutant unable to adapt rapidly in changing conditions. Because of rapid adaptability, wild-type cells utilises a major portion of the nutrient and space, leaving the Δ*ntr*C mutant deprived of nutrients (Fig.6a). Thus, in every cycle of nutrient replenishment, wild-type got significant gain over slow growing Δ*ntr*C mutant cells. This results in a drastic reduction in Δ*ntr*C mutant population in competition with wild-type, under feast-or-famine cycle. We observed an even more drastic reduction in Δ*ntr*C mutant population, when the magnitude of competition was increased by increasing the number of competitors (Fig.3f). Previous studies on competitive fitness in serial transfer to fresh medium explained that the survival is favoured by rapid exponential growth as well as by the ability to respond quickly following transfer to fresh media. Therefore, the maximum growth rate and length of lag phase following transfer to fresh medium account for competitive fitness of bacterial strains (20, 34, 35). In the first part of discussion, we have shown that Δ*ntr*C mutant has both extended lag phase and slow growth rate than the wild-type cells. Therefore, *ntr*C confer competitive fitness in both, stable and fluctuating nutrient condition, by reducing the length of lag phase and increase the growth rate.

### The lack of NtrC would lead to poor nutrient scavenging from the dead cell debris

As explained above, nutrient recycling from the dead cells is a very important strategy to maintain survival under long term starvation. Since, dying cells degrade and diffuse intracellular biomolecules, such as degradation of ribosomes, supplies ribose sugar and amino acids. Degradation of other intracellular biomolecules supplies amino acids, peptides and metal ions that can be reused by the viable cell to sustain under long-term starvation(25, 33). Therefore, the gene expression studies in dead cell extract showed that Δ*ntr*C mutant has down-regulation of genes involved in transport of amino acids such as glutamine, histidine, and arginine. Besides this, Δ*ntr*C mutant also showed down-regulation of gene encoding transporters of dipeptides, tripeptides, branched chain amino acids and metal ions. Thus, the suppressed expression of these transporter genes may be responsible for poor nutrient recycling efficiency in Δ*ntr*C mutant. While the poor nutrient recycling efficiency may be one of the factor for inability of Δ*ntr*C mutant to survive under long term nitrogen starvation and in competition with wild-type cells in stable environment.

## Conclusion

NtrC, is a global regulator and plays an important role in cell survival in low nutrient condition other than just nitrogen scavenging. NtrC protein is needed for rapid transition from lag to log phase. The NtrC provides a competitive edge by facilitating cells with a rapid adaptation system that senses and responds to the environmental changes including feast-or-famine kind of situation (Fig.8). Hence, NtrC is inevitable for an enteric pathogen *Salmonella* in a competitive environment.

**Fig.8.**
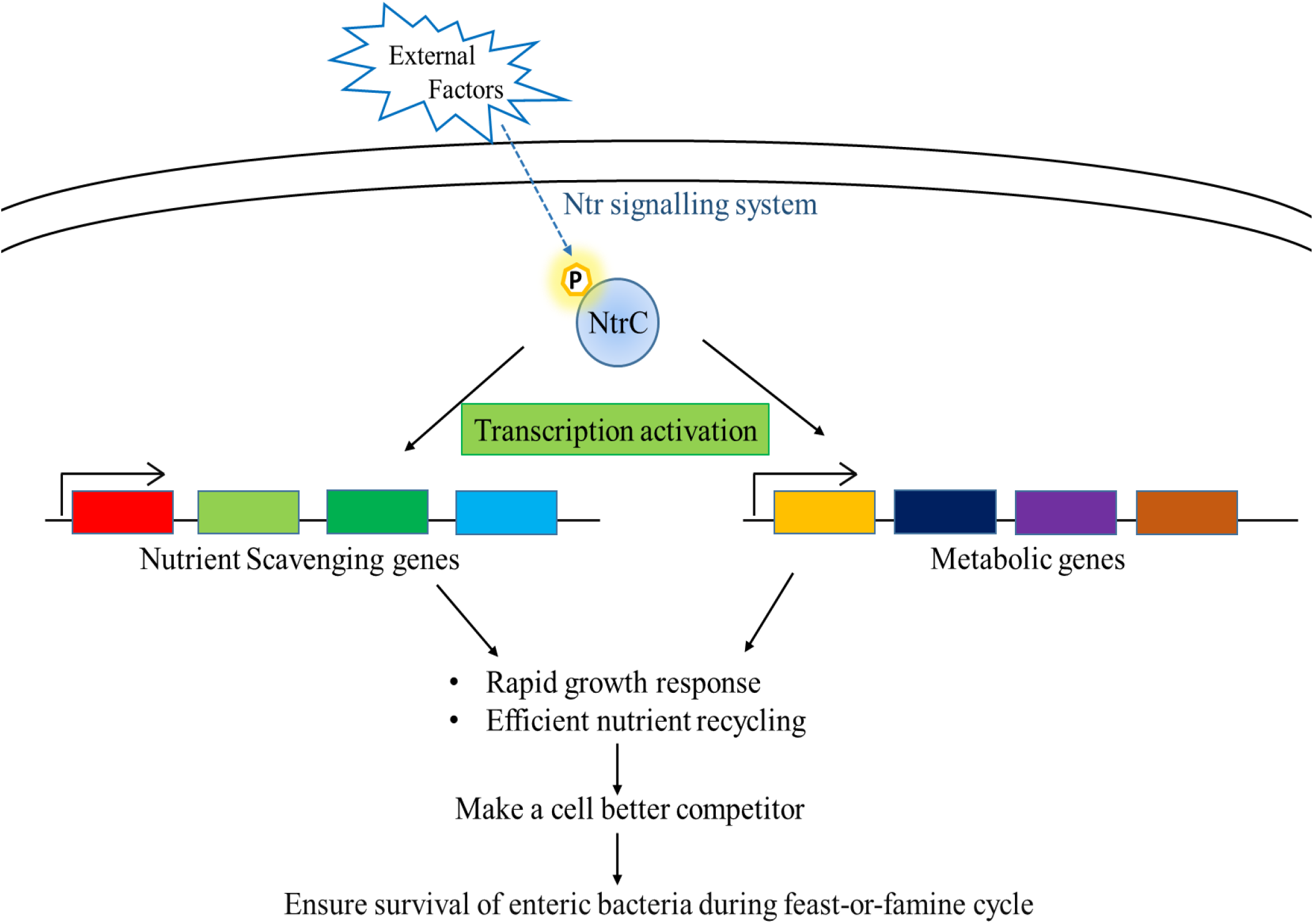
Schematic diagram on the role of NtrC in *S*. Typhimurium competitve fitness during feast-or-famine cycle. NtrC expression at the early phase of growth, provide competitve fitness and adaptation to long term nitrogen starvation, by faciliting cell with an assentail process of gene activation in response to extranal stimuli. NtrC activates genes/operons involve in nutient scavenging and metabolism. Which results in, generation of rapid growth response and efficient nutrient recycling/scavenging, that in turn confer competitive fitness under low and fluctuating nutrient condition.

## Acknowledgements

We thank the Department of Atomic Energy, Government of India, for financial support.

## reference

1. Wanjugi P, Fox GA, Harwood VJ. 2016. The Interplay Between Predation, Competition, and Nutrient Levels Influences the Survival of Escherichia coli in Aquatic Environments. Microb Ecol 72:526–37.

2. Bauer MA, Kainz K, Carmona-Gutierrez D, Madeo F. 2018. Microbial wars: Competition in ecological niches and within the microbiome. Microbial cell (Graz, Austria) 5:215–219.

3. Lam LH, Monack DM. 2014. Intraspecies competition for niches in the distal gut dictate transmission during persistent Salmonella infection. PLoS Pathog 10:e1004527.

4. Rychlik I, Barrow PA. 2005. Salmonella stress management and its relevance to behaviour during intestinal colonisation and infection. FEMS Microbiology Reviews 29:1021–1040.

5. Zimmer DP, Soupene E, Lee HL, Wendisch VF, Khodursky AB, Peter BJ, Bender RA, Kustu S. 2000. Nitrogen regulatory protein C-controlled genes of *Escherichia coli*: Scavenging as a defense against nitrogen limitation. Proceedings of the National Academy of Sciences 97:14674.

6. Santos-Beneit F. 2015. The Pho regulon: a huge regulatory network in bacteria. 6.

7. Skaar EP. 2010. The battle for iron between bacterial pathogens and their vertebrate hosts. PLoS Pathog 6:e1000949.

8. Merrick MJ, Edwards RA. 1995. Nitrogen control in bacteria. Microbiological reviews 59:604–622.

9. Yang Z, Li Q, Yan Y, Ke X, Han Y, Wu S, Lv F, Shao Y, Jiang S, Lin M, Zhang Y, Zhan Y. 2021. Master regulator NtrC controls the utilization of alternative nitrogen sources in Pseudomonas stutzeri A1501. World Journal of Microbiology and Biotechnology 37:177.

10. Hervás AB, Canosa I, Santero E. 2008. Transcriptome analysis of Pseudomonas putida in response to nitrogen availability. Journal of bacteriology 190:416–420.

11. Reitzer LJ, Magasanik B. 1983. Isolation of the nitrogen assimilation regulator NR(I), the product of the glnG gene of Escherichia coli. Proceedings of the National Academy of Sciences of the United States of America 80:5554–5558.

12. Weiss V, Claverie-Martin F, Magasanik B. 1992. Phosphorylation of nitrogen regulator I of Escherichia coli induces strong cooperative binding to DNA essential for activation of transcription. Proceedings of the National Academy of Sciences of the United States of America 89:5088–5092.

13. Weiss DS, Batut J, Klose KE, Keener J, Kustu S. 1991. The phosphorylated form of the enhancer-binding protein NTRC has an ATPase activity that is essential for activation of transcription. Cell 67:155–67.

14. Goodall ECA, Robinson A, Johnston IG, Jabbari S, Turner KA, Cunningham AF, Lund PA, Cole JA, Henderson IR. 2018. The Essential Genome of Escherichia coli K-12. mBio 9.

15. Warsi OM, Andersson DI, Dykhuizen DE. 2018. Different adaptive strategies in E. coli populations evolving under macronutrient limitation and metal ion limitation. BMC Evolutionary Biology 18:72.

16. Schink SJ, Biselli E, Ammar C, Gerland U. 2019. Death Rate of E. coli during Starvation Is Set by Maintenance Cost and Biomass Recycling. Cell Syst 9:64–73.e3.

17. Bustin SA, Benes V, Garson JA, Hellemans J, Huggett J, Kubista M, Mueller R, Nolan T, Pfaffl MW, Shipley GL, Vandesompele J, Wittwer CT. 2009. The MIQE Guidelines: Minimum Information for Publication of Quantitative Real-Time PCR Experiments. Clinical Chemistry 55:611–622.

18. Bertrand RL. 2019. Lag Phase Is a Dynamic, Organized, Adaptive, and Evolvable Period That Prepares Bacteria for Cell Division. J Bacteriol 201.

19. Hamill PG, Stevenson A, McMullan PE, Williams JP, Lewis ADR, S S, Stevenson KE, Farnsworth KD, Khroustalyova G, Takemoto JY, Quinn JP, Rapoport A, Hallsworth JE. 2020. Microbial lag phase can be indicative of, or independent from, cellular stress. Sci Rep 10:5948.

20. Smith HL. 2011. Bacterial competition in serial transfer culture. Math Biosci 229:149–59.

21. Moreno-Gámez S, Kiviet DJ, Vulin C, Schlegel S, Schlegel K, van Doorn GS, Ackermann M. 2020. Wide lag time distributions break a trade-off between reproduction and survival in bacteria. Proceedings of the National Academy of Sciences 117:18729.

22. Gray DA, Dugar G, Gamba P, Strahl H, Jonker MJ, Hamoen LW. 2019. Extreme slow growth as alternative strategy to survive deep starvation in bacteria. Nature Communications 10:890.

23. Phaiboun A, Zhang Y, Park B, Kim M. 2015. Survival kinetics of starving bacteria is biphasic and density-dependent. PLoS Comput Biol 11:e1004198.

24. Ballantyne EN. 1930. On Certain Factors Influencing the Survival of Bacteria in Water and in Saline Solutions. Journal of bacteriology 19:303–320.

25. Takano S, Pawlowska BJ, Gudelj I, Yomo T, Tsuru S. 2017. Density-Dependent Recycling Promotes the Long-Term Survival of Bacterial Populations during Periods of Starvation. mBio 8:e02336–16.

26. Chang C, Stewart RC. 1998. The Two-Component System1: Regulation of Diverse Signaling Pathways in Prokaryotes and Eukaryotes. Plant Physiology 117:723–731.

27. Nixon BT, Ronson CW, Ausubel FM. 1986. Two-component regulatory systems responsive to environmental stimuli share strongly conserved domains with the nitrogen assimilation regulatory genes ntrB and ntrC. Proceedings of the National Academy of Sciences 83:7850.

28. Rolfe MD, Rice CJ, Lucchini S, Pin C, Thompson A, Cameron AD, Alston M, Stringer MF, Betts RP, Baranyi J, Peck MW, Hinton JC. 2012. Lag phase is a distinct growth phase that prepares bacteria for exponential growth and involves transient metal accumulation. J Bacteriol 194:686–701.

29. Kim M, Zhang Z, Okano H, Yan D, Groisman A, Hwa T. 2012. Need-based activation of ammonium uptake in Escherichia coli. Mol Syst Biol 8:616.

30. Schumacher J, Behrends V, Pan Z, Brown DR, Heydenreich F, Lewis MR, Bennett MH, Razzaghi B, Komorowski M, Barahona M, Stumpf MP, Wigneshweraraj S, Bundy JG, Buck M. 2013. Nitrogen and carbon status are integrated at the transcriptional level by the nitrogen regulator NtrC in vivo. mBio 4:e00881–13.

31. Switzer A, Burchell L, McQuail J, Wigneshweraraj S. 2020. The Adaptive Response to Long-Term Nitrogen Starvation in Escherichia coli Requires the Breakdown of Allantoin. J Bacteriol 202.

32. Brown DR. 2019. Nitrogen Starvation Induces Persister Cell Formation in Escherichia coli. Journal of bacteriology 201:e00622–18.

33. Reeve CA, Bockman AT, Matin A. 1984. Role of protein degradation in the survival of carbon-starved Escherichia coli and Salmonella typhimurium. Journal of bacteriology 157:758–763.

34. Levin BR. 1972. Coexistence of two asexual strains on a single resource. Science 175:1272–4.

35. Lenski RE, Travisano M. 1994. Dynamics of adaptation and diversification: a 10,000-generation experiment with bacterial populations. Proceedings of the National Academy of Sciences 91:6808.

